# The scramblases VMP1 and TMEM41b are required for primitive endoderm specification by targeting WNT signaling

**DOI:** 10.1101/2023.10.31.564914

**Authors:** Markus Holzner, Tea Sonicki, Hugo Hunn, Federico Uliana, Weijun Jiang, Vamshidhar R. Gade, Karsten Weis, Anton Wutz, Giulio Di Minin

**Author notes:** corresponding authors (AW); (GDM).

## Abstract

The ER resident proteins VMP1 and TMEM41b share a conserved DedA domain, which confers lipid scramblase activity. Loss of either gene results in embryonic lethality in mice and defects in autophagy and lipid droplet metabolism. We set out to investigate their role in pluripotency and specification. For this purpose, we generated Vmp1 and Tmem41b mutations in mouse embryonic stem cells (ESCs). We observed that ESCs carrying mutations in Vmp1 and Tmem41b show robust self-renewal and an unperturbed pluripotent expression profile but accumulate LC3-positive autophagosomes and lipid droplets consistent with defects in autophagy and lipid metabolism. ESCs carrying combined mutations in Vmp1 and Tmem41b can differentiate into a wide range of embryonic cell types. However, differentiation into primitive endoderm-like cells in culture is impaired, and the establishment of extra- embryonic endoderm stem (XEN) cells is delayed. Mechanistically, we show the deregulation of genes that are associated with WNT signaling. This is further confirmed by cell surface proteome profiling, which identified a significant reduction of the WNT receptor FZD2 at the plasma membrane in Vmp1 and Tmem41b double mutant ESCs. Importantly, we show that transgenic expression of Fzd2 rescues XEN differentiation. Our findings identify the role of the lipid scramblases VMP1 and TMEM41b in WNT signaling during extra-embryonic endoderm development and characterize their distinct and overlapping functions.

## INTRODUCTION

Vesicle trafficking and membrane synthesis are intricately interconnected within cellular signaling networks. Lipid scramblases, with their property to shape lipid membrane composition, play a crucial role in facilitating inter-organelle communication. Beyond their integral involvement in maintaining cell functionality, recent studies have unveiled their implications in various pathological conditions, such as viral infection [1–5], cancer development [6–11], and inflammatory diseases [12–16]. The vacuole membrane protein 1 (VMP1) and transmembrane protein 41B (TMEM41B) are lipid scramblases with a highly conserved DedA domain [17, 18]. They localize at the ER membrane and are distinguished from other ER-associated flippases or scramblases by the property to facilitate bi-directional lipid diffusion in an ATP- and Ca2+-independent manner. VMP1 and TMEM41B are necessary for autophagosome formation [19, 20], lipid droplet metabolism [17, 21], and lipoprotein formation [22]. Additional reports proposed a more global role in ER protein homeostasis [23] and vesicle transport [24]. Despite multifaceted functions, their absence does not influence cell survival significantly. Compensatory mechanisms shared by VMP1 and TMEM41B or other ER scramblases likely maintain cellular homeostasis. While VMP1 depletion leads to a pronounced defect in autophagosome assembly [19, 20], TMEM41B appears to play a pivotal role in sustaining lipid droplet homeostasis and serves as a crucial factor in RNA virus replication [3, 25]. However, a direct comparison between the activities of VMP1 and TMEM41B has not yet been performed.

VMP1 and TMEM41B are both essential during embryonic development. Vmp1 [21] and Tmem41b [26] deficient mice die around E8.5 with a severe developmental delay. Previously, defects in the visceral endoderm function have been proposed. In particular, loss of Vmp1 has been suggested to block the synthesis and release of lipoproteins from the visceral endoderm. However, mice with defects in lipoprotein biogenesis experience mortality at a later stage, E10.5 [27, 28]. This observation suggests a distinct and earlier involvement of VMP1 and TMEM41B in embryo development.

During implantation, the mouse embryo is composed of three lineages: the trophectoderm, the primitive endoderm (PE), and the epiblast (EPI), which is primed to develop into the embryo [29]. The EPI and PE are specified within the inner cell mass (ICM) of the blastocyst. Defects in the formation of these lineages are challenging to characterize *in vivo* as they appear pre-implantation and due to the small embryo size. To investigate the role of Vmp1 and Tmem41b in early embryonic lineage specification, we generated mutations in Vmp1 and Tmem41b in ESCs. We show that a combined Vmp1 and Tmem41b mutation in ESCs results in a specific defect in PE formation and a delayed conversion to extra-embryonic endoderm cells (XEN). This delay is caused by reduced WNT signaling and can be rescued through WNT activation.

## RESULTS

### Vmp1 and Tmem41b mutant ESCs accumulate LC3+ autophagosomes and lipid droplets

To assess the influence of a Vmp1 or Tmem41b depletion on mouse ESCs and in early embryo development, we engineered genomic deletions using CRISPR/Cas9 targeted mutagenesis. Vmp1 was mutated by introducing a frameshift in exon 2 (Fig.1A). Loss of Vmp1 protein was confirmed in independent mutant ESC clones (Vmp1^KO^) (Fig.S1A). To abrogate Tmem41b expression, we deleted the genomic region between exons 2 and 3 (Fig.1A), which encodes the first transmembrane domain of the protein (Tmem41b^KO^). This mutation is predicted to compromise the protein topology. Additionally, the splicing of exon 1 to exon 4 produces a frameshift, suggesting a null allele. The deletion was confirmed on the genomic DNA (Fig.S1B). Tmem41b mRNA levels were strongly reduced (Fig.S1C-D).

**Figure 1.**
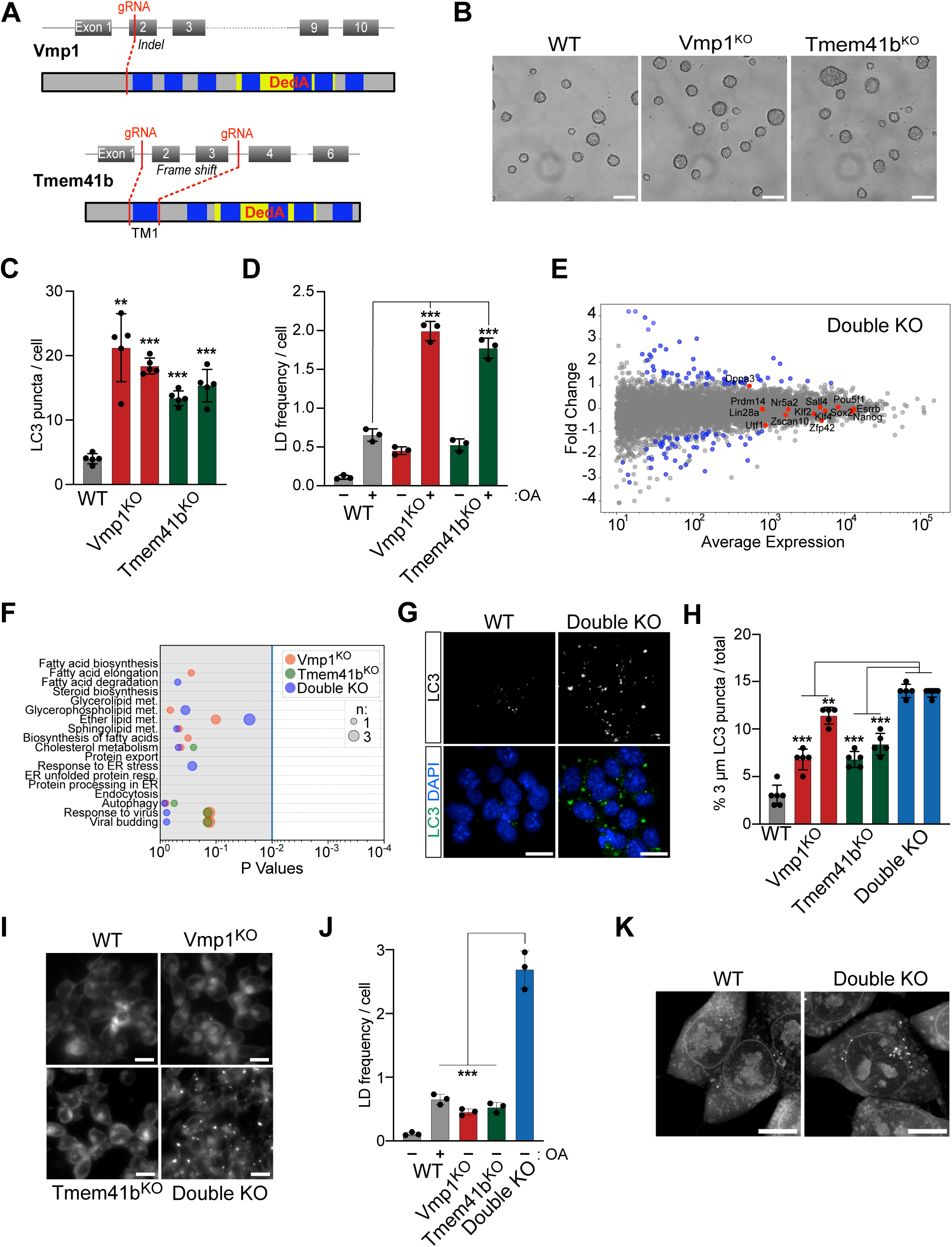
Vmp1 and Tmem41b double KO ESCs display increased autophagosome and lipid droplet accumulation. **(A)** Schematic representation of the CRISPR/Cas9-mediated KO strategy for Vmp1 and Tmem41b genes. For depleting Vmp1 (top) a single gRNA inducing an indel mutation has been used. In the case of the Tmem41b gene (bottom), two gRNAs were used to delete exons 2 and 3. The sequences coding for transmembrane domains (TM) are indicated in blue, the conserved DedA domain is highlighted in yellow, red lines indicate cut sites of gRNAs. **(B)** Colony morphology of VMP1^KO^ and Tmem41^KO^ ESCs remains unchanged compared to WT ESCs. Scale bar: 100 µm. **(C)** VMP1 and Tmem41b depletion induce accumulation of autophagosomes in ESCs. The graph shows the quantification of LC3 puncta per cell of a representative experiment (see Fig.S2A). Data were collected from more than 200 cells. For each KO condition, two independent clones have been analyzed. Dots correspond to the frames analyzed per sample. Data are represented as mean ± SD. The P-value is relative to the WT sample. **(D)** Vmp1 and Tmem41b depletion leads to the accumulation of lipid droplets (LD) in ESCs. Graph quantifies LD per cell in WT and KO ESCs (see Fig.S2D). ± oleic acid treatment. At least 500 cells were analyzed for every condition; dots represent independent biological replicates. Data are represented as mean ± SD. (**E-G)** Transcriptional analysis of Double KO ESCs did not identify many deregulated genes. **(E)** Shown are differentially regulated genes in Double KO ESCS in blue as MA plot. Genes highlighted in red are pluripotency markers. **(F)** Differentially regulated genes do not cluster in essential processes. The graph shows differentially expressed genes of single and double KO ESCs by cellular processes. Dot size correlates to the number of deregulated genes within a cellular process. **(G-H)** Double KO ESCs accumulate LC3+ autophagosomes of increased size. **(G)** Shown are representative IF images of WT and double KO ESCs stained for LC3. Nuclei are stained with Dapi. Scale bar: 20 µm. **(H)** Plotted is the quantification of LC3 puncta of 3 µm in diameter. For each condition, two independent clones have been analyzed. Dots correspond to the frames analyzed per sample. Data were collected from more than 200 cells. Data are represented as mean ± SD. The P-value is relative to the Double KO sample with the highest SD. **(I-J)** Double KO ESCs accumulate LDs without oleic acid treatment. **(I)** Live cell staining of WT, Vmp1, Tmem41b, and double KO ESCs with BODIPY. Scale bar: 20 µm **(J)** The number of LDs per cell is plotted for all KO clones. Data are represented as mean ± SD. **(K)** Double KO ESCs present an increased amount of LD visualized through refractive index microscopy (top) and co-stained with BODIPY.

Vmp1 and Tmem41b mutant ESCs showed similar morphology (Fig.1B), proliferation (Fig.S1E), and expression of pluripotency markers (Fig.S1F-G) compared to WT ESCs. We confirmed that the pluripotency network was unchanged by a transcriptomic analysis performed in individual Vmp1 and Tmem41b mutant cells. This analysis also showed that the transcription profile of cells was marginally affected, consistent with earlier results of ESC characterization (Fig.S1I-J and Table S2).

Considering the role of Vmp1 and Tmem41b in autophagy, we assessed the accumulation of LC3+ autophagosomes to determine the impact of their mutations. We found an increase of LC3+ puncta in both mutants compared to WT ESCs (Fig.1C and Fig.S2A-B). Interestingly, the increased frequency of LC3+ puncta was similar in Vmp1 (4X) and Tmem41b (3,5X) mutant cells. Next, mutant cells were analyzed for the retention of lipid droplets (LD). To amplify LDs, we treated all samples with oleic acid (OA) and visualized LDs by BODIPY staining. We observed that Vmp1 and Tme41b mutations led to a 4-fold increase of LD numbers with respect to WT cells (Fig.1D and Fig. S2D-E). These results suggest that both Vmp1 and Tmem41b contribute to autophagosome and lipid droplet metabolism in ESCs, recapitulating earlier studies performed in other cellular systems [17, 20]. The direct comparison of both mutations revealed a similar contribution of VMP1 and TMEM41B to both processes.

### Vmp1 and Tmem41b activity is not required for ESC self-renewal

We generated double mutant cell lines to characterize potential redundant functions between VMP1 and TMEM41B. A Tmem41b mutation was engineered in Vmp1 mutant ESCs (Double KO) (Fig.S1A-D). Several independent mutant clones were obtained. Notably, Double KO cells showed a regular ESC morphology (Fig.S1H) and proliferation rate (Fig.S1E). The absence of Vmp1 and Tmem41b did not affect secretory compartments, as ER, Golgi apparatus, and late endosomes appeared normal in mutant cells (Fig.S3A). Transcriptomics analysis of Double KO cells showed a limited perturbation in gene expression (Fig.1E). The combined mutations did not result in a greater number of differentially regulated genes compared to Vmp1^KO^ cells (Fig.S3C). Minimal overlap of differentially expressed genes was detected among Vmp1, Tmem41b, and Double KO mutant cell lines (Fig.S3B-C). Additionally, we did not observe enrichment for genes associated with defined cellular processes (Fig.1F, Fig.S3D-E). The pluripotency network, as also shown by IF staining (Fig.S1F-G), was not perturbed in Double KO cells. We conclude that Vmp1 and Tmem41b are not essential for ESC self-renewal.

### Vmp1 and Tmem41b have redundant functions in autophagosome and lipid droplet formations

LC3 staining showed that Double KO cell lines present an exacerbated phenotype in autophagosome accumulation compared to single mutants. Although the total number of LC3 positive autophagosomes per cell remained unchanged (Fig.1G and Fig.S2A- B), we observed an increased size. 14% of LC3 clusters in the Double KO had a larger size (>3μm), compared to the 8% of autophagosomes across the single mutations and 3% in the WT condition (Fig.1H and Fig.S2C).

The simultaneous depletion of Vmp1 and Tmem41b also had a greater impact on LD formation. Double KO cells showed a drastic increase of LD number in the absence of oleic acid treatment, whereas in WT and Vmp1 and Tmem41b mutant ESCs, LDs became detectable after OA treatment (Fig.1I-J and Fig.S2D-E). As BODIPY could also stain other lipid-containing compartments, including autophagosomes [30], we confirmed the enriched lipid droplets in Double KO cells by refractive index tomography, which is a labelling-independent technique (Fig.1K and Fig.S2F-G). Together, these results indicate that Vmp1 and Tmem41b have overlapping functions in ESCs, as the combined mutation of both genes exacerbates autophagosome and lipid droplet accumulation.

### Loss of Vmp1 and Tmem41b affects ESC potential to differentiate into primitive endoderm

To evaluate the effect of Vmp1 and Tmem41b on the differentiation potential of ESCs, we used embryoid body (EB) differentiation (Fig.2A). After 6 days of differentiation, EBs derived from WT ESCs showed increased expression of ectoderm (Sox1 and Fgf5), mesoderm (Actc1 and Brachyury) and endoderm (Gata6 and Dab2) genes (Fig.2B and Fig.S4A), while the expression of pluripotency markers (Nanog and Pou5f1) was lost (Fig.2B and Fig.S4A). We evaluated variations in the differentiation potential of mutant cells by comparing the expression of these markers by RT-qPCR. For each mutant, the differentiation potential of two independent clones was tested. Vmp1, Tmem41b, and double mutant ESCs formed well-defined EBs that resembled those of WT cells. All cell lines efficiently differentiated into the ectoderm and mesoderm lineages (Fig.2C and Fig.S4B-C). Notably, Double KO EBs showed a decreased expression of the endoderm markers Dab2 (100%) and Gata6 (57% of experiments) (Fig.2C and Fig.S4C). Neither single mutant showed a similar failure in Dab2 expression, even if some variability was detected between the analyzed clones. In the single mutants, we noticed a decreased endoderm specification, as shown by Gata6 expression levels in experiments performed with Vmp1^KO^ (56% of experiments) and Tmem41b^KO^ (58%) cells (Fig.2C and Fig.S4C). Our results indicate that combined mutation of Vmp1 and Tmem41b leads to reduced Dab2 expression and predisposes to variability in the expression of other endodermal genes.

**Figure 2.**
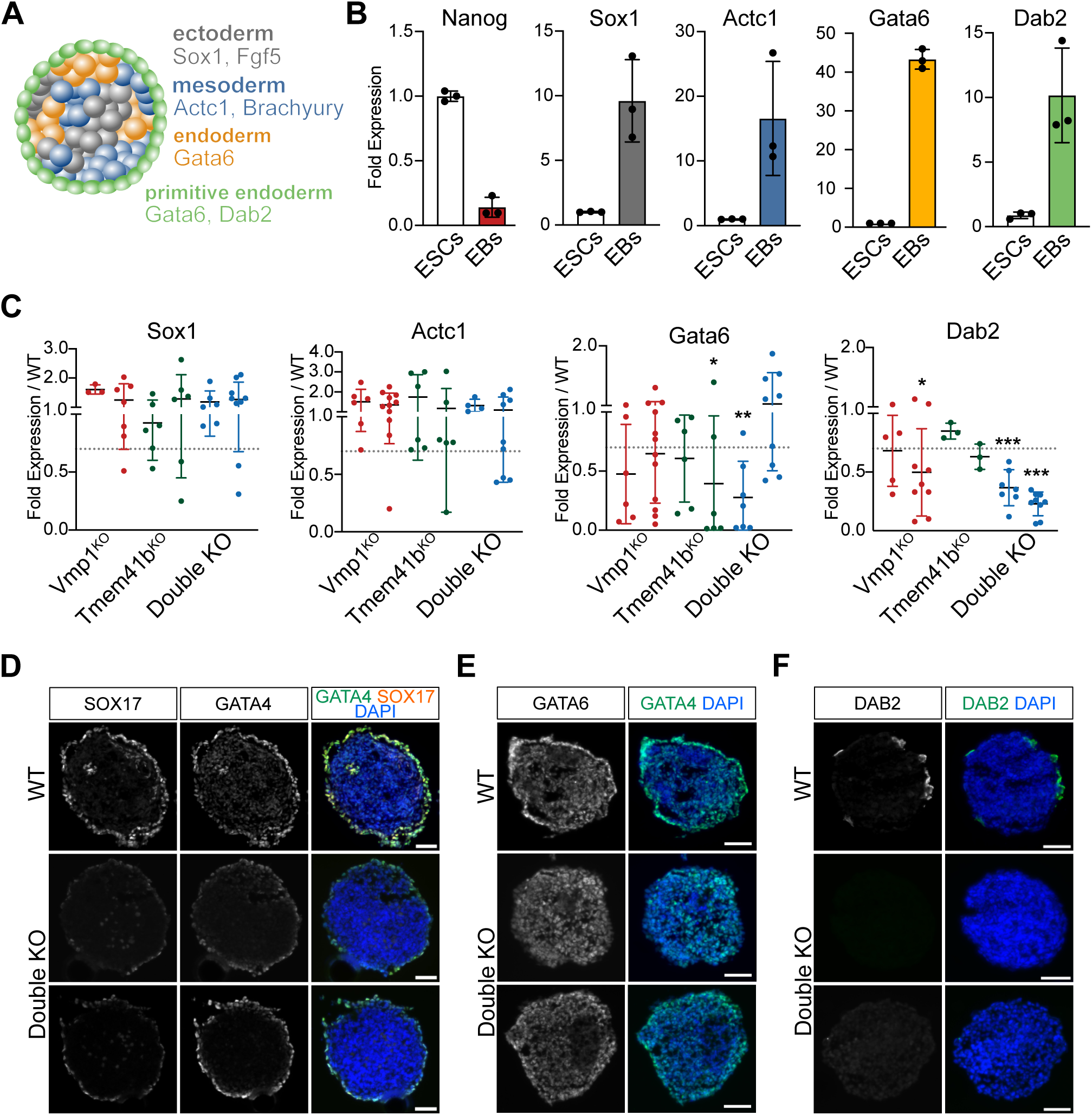
Double KO ESCs specify all germ layers, but not the primitive endoderm. **(A)** Schematic representation of an Embryonic Body (EB) which consists of 3 different lineages. The ectoderm expresses Fgf5 and Sox1 (grey), the mesoderm Actc1 and Brachyury (blue), and the endoderm by Gata6. The surrounding region of the EB corresponds to the primitive endoderm (green) and is marked by Gata6 and Dab2 expression. **(B)** ESCs differentiated to EBs express lineage markers for ectoderm (Fgf5), mesoderm (Brachyury), endoderm (Gata6), and primitive endoderm (Dab2). Graphs show mRNA levels of WT EBs normalized to the Sdha gene. Dots represent biological independent experiments. Data are represented as mean ± SD. **(C)** Double KO ESCs specifically show defects in expressing the primitive endoderm marker, Dab2. qPCR analysis for Sox1, Actc1, and Dab2 mRNA expression in mutant EBs relative to WT EBs. Two independent clones were analyzed per cell line. Each dot represents an independent experiment. Data are represented as mean ± SD. The line at 0,7 indicates an arbitrary threshold after which successful expression of the marker is considered. Statistical significance (Welch’s t test) is denoted as follows: ns: p> 0.05, *: p < 0.05, **: p < 0.01, ***: p < 0.001. **(D-F)** Double mutant EBs show decreased expression of endoderm markers. Immunofluorescence images of EB sections stained for the endoderm markers GATA4 and SOX17 **(D)**, GATA6 **(E)**, DAB2 **(F)**. WT in the top panel, two independent clones for Double KO condition in the center and bottom panels. Nuclei are stained with Dapi. Scale bar: 50 µm.

We confirmed the differentiation potential of Double KO cells into ectodermal lineages by generating SOX1-positive spinal cord organoids (Fig.S4D). Mesodermal differentiation was inferred from the ability of double mutant EBs to form contractile cardiomyocytes after plating (Video S1-2). We subsequently sectioned day 6 EBs to evaluate protein expression and distribution of endoderm markers by IF. Expression of GATA4, GATA6, and SOX17 was detected in all conditions but noticeably decreased in Double KO compared to WT EBs (Fig.2D-E). Interestingly, decreased GATA4 and GATA6 staining was pronounced in EB border regions, which resemble the primitive endoderm layer [31]. In line with these findings, we confirmed the absence of DAB2 in Double KO EBs (Fig.2F). DAB2 is specifically expressed in the primitive endoderm, while GATA4 and GATA6 are known to be also expressed in definitive endoderm.

The significant reduction in both mRNA and protein levels of Dab2, along with the weakened expression of endoderm markers at the edges of embryoid bodies, provides evidence that the absence of Vmp1 and Tmem41b compromises the formation of primitive endoderm.

### Combined Vmp1 and Tmem41b mutations delay ESC differentiation into extra- embryonic endoderm stem (XEN) cells

We tested the ability of Double KO ESCs to differentiate into extra-embryonic endoderm stem (XEN) cells to characterize endodermal differentiation further. XEN cells share characteristics with PE cells including cell signaling, gene expression, and differentiation potential [32]. Double KO and WT cells were differentiated into XEN cells with Activin A and Retinoic Acid [33]. After 6 days, WT ESCs efficiently differentiated into XEN clusters (Fig.3A), exhibiting a stellate and refractile morphology [33]. In Double KO cells, the formation of XEN clusters was substantially reduced. We characterized XEN identity by analyzing the expression of XEN markers by RT-qPCR (Fig.S5A) and immunostaining (Fig.3B and Fig.S5B). It is worth noting that the small number of derived XEN cells from Double KO ESCs can be expanded for multiple passages, as shown in Fig.S5C. This result suggests that the loss of Vmp1 and Tmem41b does not entirely prevent XEN specification, but rather causes a delay in the differentiation process.

**Figure 3.**
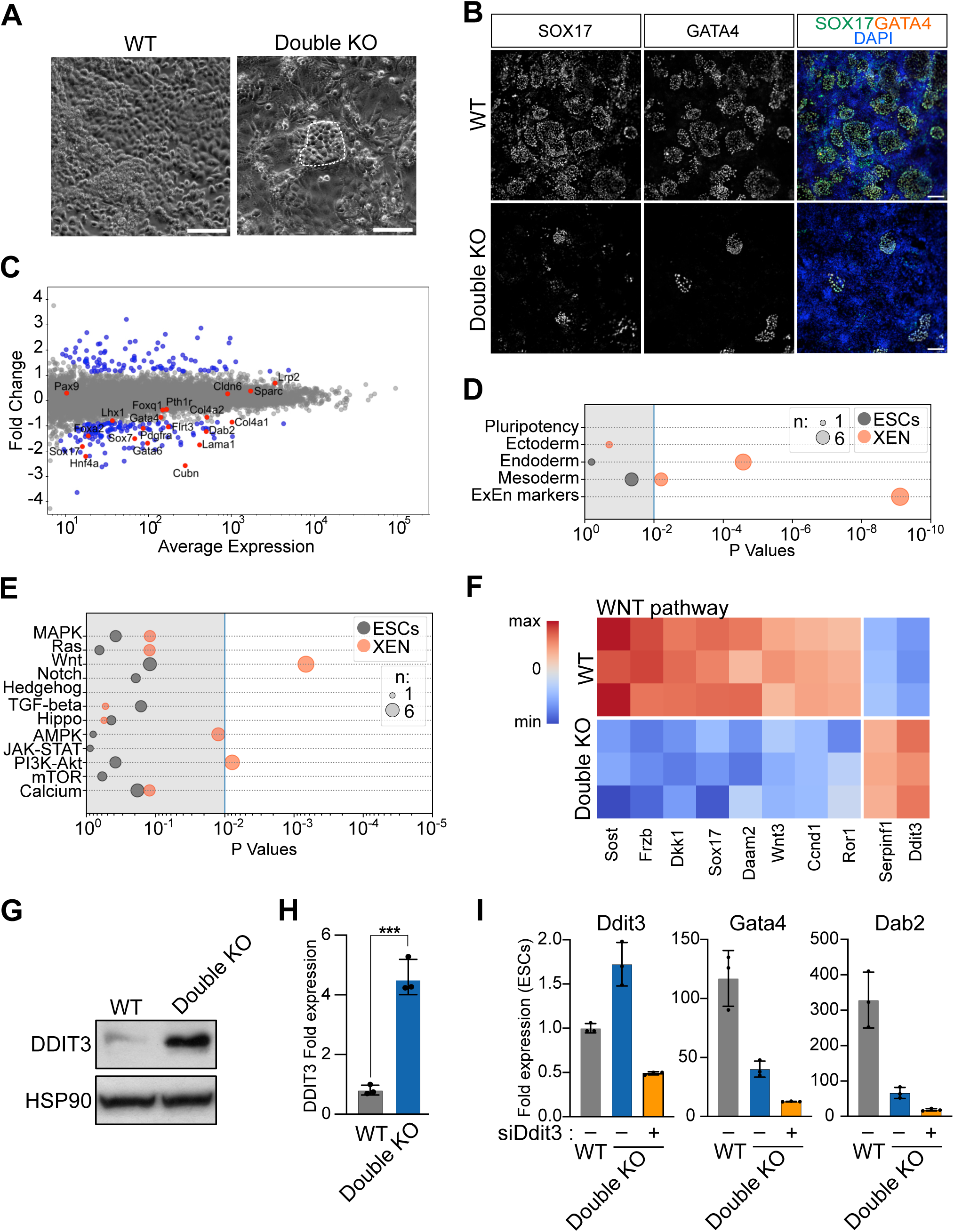
XEN differentiation in Double KO cells is delayed (A-B) Double KO ESCs show a delay in XEN differentiation. **(A)** Brightfield images of XEN cells at day 6 of differentiation in WT and Double KO cells. XEN islands are circled in white in the Double KO condition. Scale bar: 100 µm. **(B)** Immunofluorescence images of day 6 XEN cells stained for SOX17 and GATA4. Nuclei are stained with Dapi, Scale bar: 150 µm. **(C-F)** Transcriptional analysis of XEN cells. **(C)** Differentially expressed genes between Double KO and WT cells at day 2 during XEN specification. XEN markers are labeled in red; differentially expressed genes in blue. **(D-E)** GO-Term analysis of differentially expressed genes between Double KO and WT conditions in ESCs and XEN cells in the context of developmental processes **(D)** and signaling pathways **(E)**. Dot size corresponds to the number of differentially regulated genes. **(F)** WNT pathway-related genes differentially expressed in Double KO cells. Heatmap shows min to max expression of the average of three biological replicates for WT and Double KO XEN cells. **(G-H)** VMP1 and Tmem41b depletion promotes the increase of DDIT3 protein levels. **(G)** Western Blot analysis of DDIT3 expression in WT and Double KO cells differentiated to XEN cells for 2 days. HSP90 is shown as loading control. **(H)** Quantification of increased DDIT3 protein levels in three biological replicates. Expression relative to WT XEN cell and normalized to respective HSP90 expression. Data are represented as mean ± SD. **(I)** DDIT3 silencing does not restore XEN specification in Double KO cells. qPCR analysis at day 3 of differentiation for Ddit3, Gata4, and Dab2 mRNA expression in mutant cells relative to WT ESCs transfected without and with Ddit3 siRNA. Data are represented as mean ± SD (n=3).

### WNT signaling is decreased in Vmp1 and Tmem41b mutant cells during XEN specification

To characterize the effect of Vmp1 and Tmem41b double mutation on XEN specification, we analyzed the transcriptomic profile of Double KO and WT cells during the differentiation process. On day two of differentiation, we detected the induction of XEN-associated genes [34] in WT cells. However, in Double KO cells these genes were weakly expressed (Fig.3C-D and Fig.S5E). Intriguingly, we did not observe any perturbation in the transcription of genes involved in lipid metabolism, autophagy, or ER maintenance (Fig.S5D).

Instead, Double KO cells show consistent deregulation of genes associated to the WNT pathway (Fig.3E-F). A close analysis of these genes allowed to identify several well-established WNT-target genes, including Dkk, Frzb, Sox17, Wnt3, and Ccdnd1 [35–38]. Their down-regulation strongly suggested a decreased activity of the WNT pathway in Double KO cells and might explain the observed defect in XEN specification [39]. Interestingly, Ddit3 was strongly upregulated in Double KO cells. DDIT3 acts as an inhibitor of the WNT signaling by repressing the transcriptional activity of the TCF7/TCF4 complex [40]. We confirmed increased levels of the DDIT3 protein in Double KO cells by Western analysis (Fig.3G) and quantified a fourfold increase over WT cells (Fig.3H). Considering that Ddit3 is not reported to be a WNT-targeted gene, but instead, its expression is controlled by VMP1 levels [41], we verified if DDIT3 increase might explain WNT-target gene repression in mutant cells. For this purpose, we depleted Ddit3 in WT and mutant cells by RNA interference. After two days of differentiation, the transcription of XEN-specific genes was evaluated by qPCR. Despite efficient siRNA-mediated reduction of Didt3 mRNA levels (Fig. 3I), we did not observe an increase in the expression of XEN marker genes in Double KO cells.

These results indicate that the increased level of DDIT3 in mutant cells is not the only cause of the XEN specification defect caused by Vmp1 and Tmem41b mutations.

### Vmp1 and Tmem41b mutations reduce FZD2 abundance at the plasma membrane

To identify potentially affected factors of cell signaling, we investigated the influence of the Vmp1 mutation on proteins at the plasma membrane (PM) in ESCs. For this purpose, we enriched surface proteins in mutant and WT cells using aminooxy-biotinylation and characterized their abundance by TMT mass spectrometry (Fig.S6A-B). Our analysis revealed that a limited subset of proteins showed differential abundance at the plasma membrane in mutant cells, with 26 proteins significantly down-regulated and 12 up-regulated (out of 1417 detected proteins) (Fig. 4A and Table S4). This finding aligns with our RNA expression profiles, indicating that Vmp1 and Tmem41b mutations has a limited effect on the protein levels of ESCs. It also supports the conclusion that, despite defects in autophagy and lipid droplet formation, the secretory pathway remains largely intact in the absence of Vmp1. GO-term analysis highlighted that proteins more abundantly present at the PM in Vmp1 mutant cells are characterized by an ER-Golgi localization, suggesting partial defects in retrograde trafficking (Fig. 4B). Moreover, we noticed that the abundance of four GPCRs was reduced at the plasma membrane of mutant cells (Fig. 4B). Among them, we found the WNT-receptor FZD2 (Fig. 4A and Table S4). Other Frizzled receptors—FZD5, FZD7, and FZD10— were detected at the PM but remained un-affected (Fig. 4C and S6C). The decrease of FZD2 protein abundance at the PM was independent of Fzd2 transcription, as no difference in Fzd2 mRNA was detected between WT and mutant cells (Fig. S6D).

**Figure 4.**
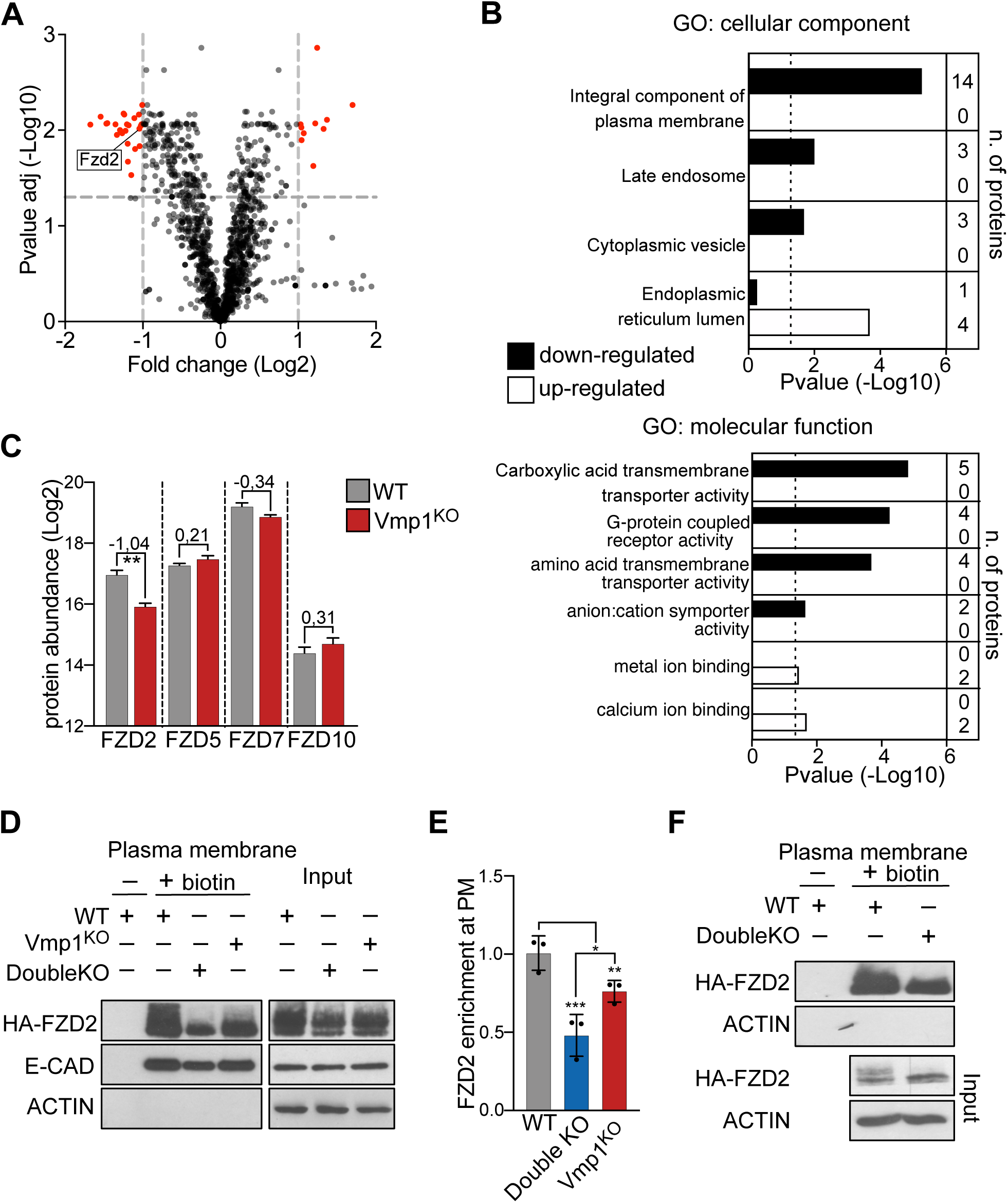
Vmp1 and Tmem41b mutant ESCs show reduced plasma membrane abundance of the WNT receptor FZD2. **(A)** Differences in the abundance of plasma membrane proteins in WT and VMP1^KO^ cells. Volcano plot displaying differentially expressed proteins at the plasma membrane in WT and VMP1^KO^ ESCs. Proteins with the most significant divergence are highlighted in red. Gray lines denote the selection criteria: an absolute log2 fold-change > 1 and a false discovery rate < 0.05. The position of FZD2 is annotated. **(B)** GO term analysis of proteins differentially abundant at the plasma membrane of WT and Vmp1^KO^ cells. Shown are GO terms related to protein localization and molecular function. **(C)** Abundance of Frizzled receptors identified by MS analysis in WT and mutant cells. The average and standard error is reported for three independent biological replicates **(D)** Western blot analysis of HA-FZD2 abundance at the PM in WT, VMP1^KO^, and Double KO ESCs. E-CADHERIN and ACTIN serve as controls for PM purification and loading, respectively. **(E)** Quantification of HA-FZD2 protein levels at the PM across three biological replicates relative to WT cells and normalized to the total HA-FZD2 (Input). Data are represented as mean ± SD. **(F)** Western blot analysis of HA-FZD2 abundance at the PM in WT and Double KO cells differentiated into XEN cells for 2 days.

To confirm these findings and assess whether FZD2 reduction contributes to the XEN specification defects in Double KO cells, we generated ESCs with constitutive expression of HA-Fzd2 across Vmp1KO, Double KO, and WT genotypes. We evaluated HA-FZD2 translocation to the plasma membrane using a cell-impermeable biotinylation reagent followed by purification of biotinylated proteins (Fig. 4D). We confirmed that FZD2 is less abundant at the plasma membrane in Vmp1 mutant cells, validating previous MS results (Fig. 4D-E). Notably, Double KO cells exhibited an even greater reduction in FZD2 levels at the PM, suggesting an additive effect of the Vmp1 and Tmem41b mutations. Similar results were observed when analyzing HA-FZD2 enrichment at the plasma membrane in Double KO and WT cells during XEN specification (Fig. 4F). In this experiment, we noticed a marked reduction of the upper form of the FZD2 doublets in mutant cells. This suggests that FZD2 is subjected to different post-translational modifications in WT and mutant cells.

### FZD2 over-expression rescues XEN differentiation defects in Vmp1 and Tmem41b mutant cells

Our observation of deregulated WNT target genes and a reduced abundance of FZD2 at the PM indicated that WNT signaling defects might impair XEN specification. To investigate if indeed the WNT signaling cascade is affected, we quantified β-CATENIN, down-stream protein, and major signal transducer of the WNT pathway. We observed a marked decrease of β-CATENIN protein in Double KO compared to WT cells (Fig. 5A). This difference was even stronger, specifically analyzing the cytoplasmic fraction of β-CATENIN that is not associated with cadherins. To explore if the defect in XEN differentiation of double mutant cells can be rescued by chemically activating WNT signaling, we employed the GSK3 kinase inhibitor (CHIR) to prevent β-CATENIN degradation. GSK3 inhibition had no effect on XEN specification in WT ESCs. However, GSK3 inhibition significantly enhanced XEN cell cluster formation in double mutant ESCs (Fig. 5B and S7A-B). Finally, to investigate whether also overexpression of FZD2 would rescue XEN differentiation defects in Double KO cells, we generated subclones with high FZD2 expression for both Double KO and WT ESCs (Fig. S7C). The increased amount of FZD2 alone did not further stimulate the WNT pathway or XEN specification in WT cells (Fig. 5C-D and S7D-F). However, FZD2 overexpression in Double KO cells was sufficient to restore their potential for differentiation into XEN cells. Collectively, our data implicate Vmp1 and Tmem41b in WNT signaling during primitive endoderm specification.

**Figure 5.**
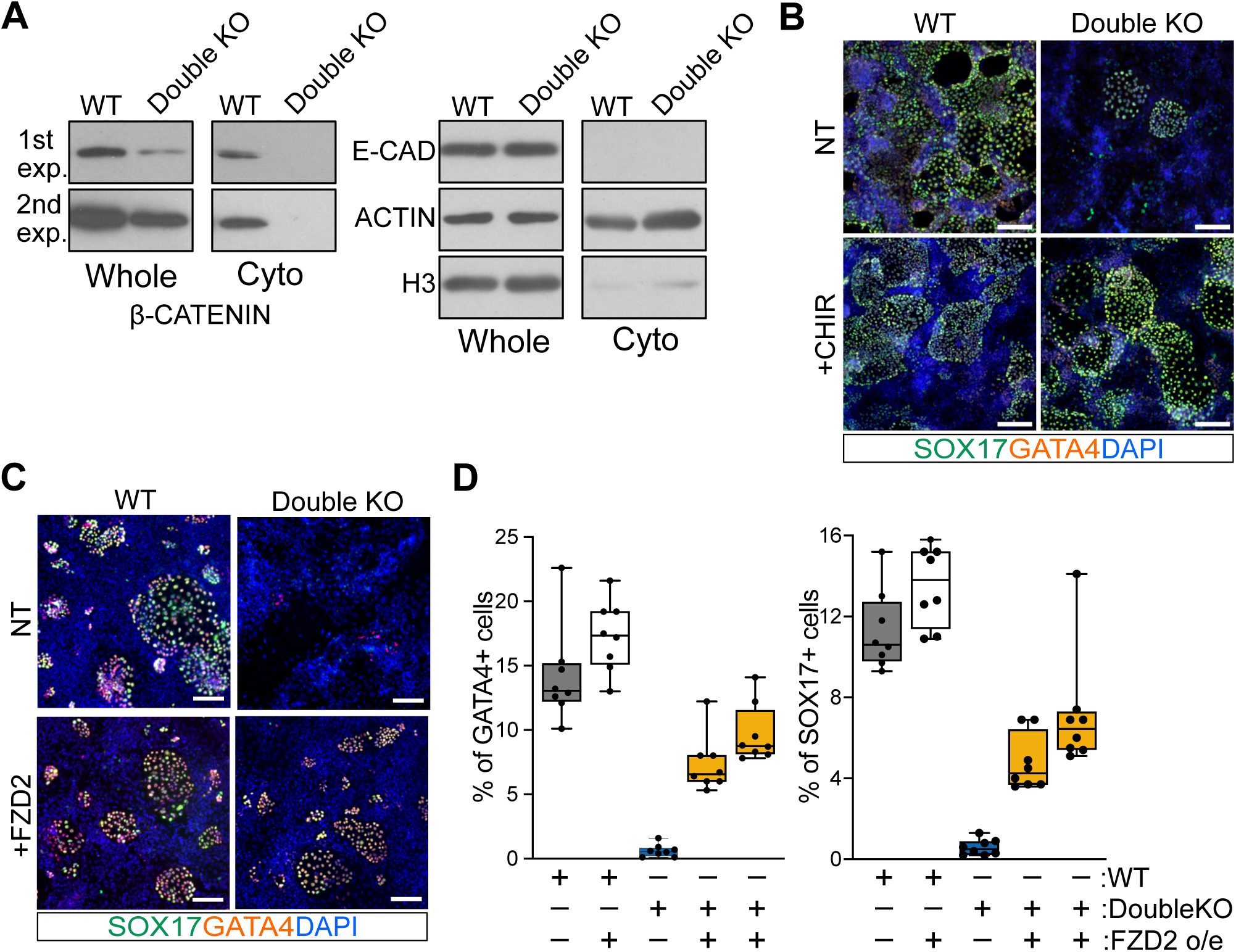
Activation of WNT signaling rescues the delay of Double KO cells in XEN differentiation. **(A)** Cytoplasmic β-CATENIN levels are reduced in Double KO cells. Western blot analysis was performed on whole-cell lysates (1/50) and purified cytoplasmic fractions (1/50) from WT and Double KO cells. E-CADHERIN, Histone H3, and ACTIN were used as controls for purification and loading. Whole-cell and purified cytoplasmic fractions are shown with the same exposure conditions. **(B)** Chiron-dependent WNT activation rescues the delay of Double KO XEN differentiation. IF staining for SOX17 and GATA4; nuclei are stained with Dapi. Scale Bar: 150 µm. **(C-D)** FZD2 overexpression rescues the differentiation delay in Double KO XEN cells. **(C)** Immunofluorescence staining for SOX17 and GATA4 in WT and Double KO cells. Nuclei are counterstained with DAPI. Scale bar: 100 µm. **(D)** Quantification of GATA4 and SOX17 positive cells during XEN differentiation, expressed as a percentage of total cells (DAPI-stained).

## DISCUSSION

Vmp1 and Tmem41b have recently been identified in a number of genetic screens designed to study various cellular processes, including autophagy, lipid metabolism, virus maturation, and ER stress [22]. In this study, we have unveiled the role of Vmp1 and Tmem41b in regulating protein secretion. Our findings indicate that their presence is essential for the trafficking of the WNT receptor FZD2 to the plasma membrane. A drastic reduction of FZD2 in mutant cells impairs WNT signals and causes an XEN cell differentiation defect. Our data, therefore, implicate a function of Vmp1/Tmem41b in WNT signaling. Given that Vmp1 expression and activity are influenced by various stimuli [42, 43], the interaction between Vmp1 and Tmem41b within the secretory pathway is critical for physiological and pathological processes.

The pleiotropic function and peculiar structure of the DedA domain found in both scramblases have prompted multiple studies addressing its mode of action and role in cellular homeostasis. An important point under evaluation is whether Vmp1 and Tmem41b work cooperatively or show specialization for different cellular processes. To address this question, we targeted Vmp1 and Tmem41b genes in ESCs and introduced mutations abrogating their expression. This allowed us to directly compare the specificity of Vmp1 and Tmem41b in multiple processes. We report that their absence similarly affects autophagy and lipid droplet accumulation. Moreover, we show that the simultaneous loss of both genes synergizes and exacerbates defects of single mutants, particularly in LD formation. Notably, loss of both genes did not affect self-renewal in ESCs and had only subtle effects on gene expression. Our characterization shows that Vmp1 and Tmem41b perform distinct and overlapping functions in ESCs. It will be interesting to understand if variating expression levels of Vmp1 and Tmem41b can account for their redundancy or dominance in different cellular systems.

The embryonic lethality caused by Vmp1 and Tmem41b mutations [21, 26] during development may be explained by the defective primitive endoderm specification that we detected *in vitro*. Embryoid body experiments and the direct differentiation of ESCs to extra-embryonic endoderm stem (XEN) cells support a specific delay in PE formation in double mutant cells. Interestingly, it has been reported that other mutations leading to PE delay cause retardation and embryo lethality at E8.5 [44], mirroring phenotypes of Vmp1 or Tmem41b mutant embryos. While both scramblases are needed in mice for regular development, the depletion of both genes is required to cause a differentiation delay in ESCs. More permissive *in vitro* growth conditions may explain the requirement for the depletion of both genes to manifest this phenotype in cell culture. It is worth noting that single Vmp1^KO^ or Tmem41b^KO^ cells also showed subtle defects in endoderm specification in embryoid bodies, albeit less consistently than Double KO cells (Fig.2C). The finding of WNT signaling deregulation in Vmp1 and Tmem41b XEN cells explains the specification delay detected in mutant cells. This is consistent with earlier reports that the WNT/TCF7L1 transcriptional response is necessary *in vitro* and *in vivo* for PE expansion [39]. Retardation – not impairment – of PE specification induced by Vmp1 or Tmem41b mutations may permit embryo implantation but the accumulation of defects likely blocks development during gastrulation.

Overexpression of the FZD2 receptor rescues the defects of Vmp1 and Tmem41b double mutant cells during XEN specification. We propose that the decreased availability of FZD2 at the plasma membrane in mutant cells causes a delay in WNT signaling activation and PE defects. The observation that Fzd2 mutant mice exhibit developmental defects at later stages [45] suggests the possibility of compensatory mechanisms among different Frizzled receptors *in vivo*. Future studies will focus on the deep characterization of other potential GPCRs subjected to Vmp1/Teme41b regulation. While our results indicate that the increased expression of the WNT repressor DDIT3 alone does not fully account for the delay in XEN specification, it may still contribute to this delay during embryonic development. The elevated levels of DDIT3 in mutant cells might exert an additive effect, further reducing WNT pathway activity.

Vmp1 and Tmem41b emerge as key factors in the secretary machinery of cells. Beyond their established roles in lipid and membrane metabolism, we demonstrate their involvement in regulating protein trafficking at the plasma membrane. Notably, this effect is transcriptional independent, as evidenced by expression analyses and experiments in cells with constitutive FZD2 expression. Furthermore, our findings indicate that Vmp1/Tmem41b depletion does not cause a global disruption of protein secretion but shows a specificity for some sub-classes of secreted proteins, such as GPCRs. In denaturing conditions, the upper forms of the FZD2 protein are less abundant in mutant cells. This finding suggests potential post-translational modification differences between the two conditions. GPCRs are tightly regulated at the ER-Golgi interface, where proper folding is essential for their route in the secretory pathway. In the Golgi, GPCRs undergo extensive glycosylation, which alters their molecular weight and activity. These results suggest a role of Vmp1/Tmem41b in promoting the maturation and stability of a specific class of transmembrane proteins.

The cellular systems generated in our study will help clarify the mechanisms shared by these scramblases in controlling protein secretion. Furthermore, the role of Vmp1 and Tmem41b in modulating the WNT cascade may extend to other signaling networks, revealing other cross-connections between cellular pathways and metabolic processes.

## MATERIALS AND METHODS

### Cell Culture

Mouse ES cells were maintained as previously described in chemically defined 2i medium with added leukemia inhibitory factor (LIF) as described in Di Minin et al. [46]. Base media composition: DMEM (Gibco, #41965039), 15% FBS (Biowest, #S1810- 500) 1% sodium pyruvate (Gibco, 11360-070), 1% non-essential amino acids (Gibco, #11140-035). Lif (2000 U/mL, homemade), PD0325901 (1 µM, AxonMedchem, #391210-10-9) and Chiron (3 µM, AxonMedchem, #252917-06-9) are added fresh before use.

### EB differentiation

To obtain embryonic bodies (EBs), 200k ESCs were plated on Day1 in ESCs base media without 2i and LIF (see cell culture) into a well of a Sphericalplate5D (Kugelmeiers). On Day4, EBs have formed and are transferred to a pHEMA coated 6cm culture dish and cultured until Day6. EBs are then collected and subjected to RNA extraction and RT-qPCR analysis or prepared for sectioning and immunofluorescence. In brief, EBs are fixed in 4% PFA for 30 min, cryoprotected in 10%, 20% and 30% sucrose and then embedded and frozen in Tissue Freezing Medium (Leica, #14020108926). 15 µm sections were prepared on a cryotome, mounted onto Superfrost Plus Adhesion slides (VWR international GmBH, #631-0108) and processed for immunofluorescence staining.

### Cardiomyocyte differentiation

EBs were formed as previously described. On Day4, EBs were collected and seeded onto a non-treated 10cm dish, maintaining ESC base media. Cardiomyocytes randomly specified and were observed as early as on Day6 of differentiation.

### Spinal Cord Organoids (SCOs) formation

For derivation of SCOs, ESCs were plated in EBs media (DMEM- F12 (Gibco) and Neurobasal Medium (Gibco) (1:1 ratio) supplemented with 200 mM L-glutamine and 10% of Knockout serum replacement) on Sphericalplate 5D dishes (Kugelmeiers AG) {Holzner, 2024 #139}. On day 2, SCOs were transferred in 10 cm tissue culture dishes and treated with RA (10 nM). SCOs were collected and processed for analysis at day 5.

### Xen Differentiation

XEN cells were obtained following a protocol by Niakan et al. with minor changes [33]. In brief, 230k cells were plated in XEN medium on a gelatin-coated 3cm dish. The next day, the media was replaced with Xen medium containing 0.1 µM retinoic acid (Thermo Fischer, #17110052) and 10 ng/mL Activin A (Peprotech, #120-14). On Day4, the medium was again replaced by XEN medium, no further supplements were added, and on Day6 samples were either analyzed by IF or qPCR.

Rescue of double KO Xen differentiation was performed by adding 1 µM Chiron (AxonMedchem, #252917-06-9) at Day2, together with the retinoic acid and Activin A. Media was changed on day 4 to Xen medium. XEN medium composition: RPMI 1640 (Gibco, #61870010), GlutaMAX^TM^ (Gibco, #61870036), 15% FBS (Biowest, #S1810-500), 0,1 mM ß-mercaptoethanol (Sigma, #BCCB9882).

### CRISPR-Cas9 editing to generate mutant ESC lines

Vmp1: the gRNA (gagacgcatagcaatgagta) was cloned into a px330 vector (Addgene, #158973). Per 3cm well, 300k ESCs were plated and transfected with 2.5 µg of gRNA containing plasmid, as well as 0.25 µg of a GFP expressing plasmid (pCMV-GFP). GFP-positive cells were enriched by FACS and plated as single cells. Colonies were picked. Analysis by WB was used to identify identified VMP1 KO lines. Tmem41b: the gRNAs (gtatgtttgacctgggcgaa and gagtgacatgtggaaatcag) were cloned into px330 vectors, and KO clones were derived as described above. PCR analysis of the edited locus (FW: atcttggacagggggagttc RV: gaccaggggttcattgtcatt and FW: cagcacacacctttaatccagc RV: gccacatagcaagcttaagagc) identified KO clones. qPCR of mRNA from Exon 2 (edited locus, N-terminal end) and Exon 7 (C-terminal end) further confirmed the absence of mRNA.

### Transfection and stable cell line generation

For derivation of *Fzd2*-HA ESCs the PB-HA-Fzd2-IRES-Neo plasmid [46] was used. Stable expression of the construct was achieved in WT, Vmp1^KO^, and DoubleKO ESCs by co-transfection of the Piggybac and the PBase plasmids. Cells were plated by limiting dilution and integration events selected with G418. Multiple clones characterized by different HA-FZD2 expressions were expanded. For the dowunregulation of Ddit3 the following siRNA were used. Cells were transfected with Lipofectamine™ RNAiMAX Transfection Reagent.

### Proliferation assay

100k ESCs were plated in a well of a MW12. 48h later, cells were collected and counted. Proliferation rates were calculated as relative factor of cells collected over cells seeded.

### LC3 staining and size characterization

100k cells were plated on cover-slips, pre-treated with Matrigel (Corning, # 354277), in N2B27 media. N2B27 composition: 50 mL Neurobasal media (Gibco, # 2348949), 50 mL Advanced DMEM F12 (Gibco, # 12634010), 4% BSA (Gibco, # 15260), 1% L-Glutamine (Gibco, # 25030024), 1% B27 supplement (Gibco, # 2336837), 0,5% N2 supplement (Gibco, # 2328253), and 0,1 mM ß-mercapotethanol (Sigma, #BCCB9882). The next day, samples were fixed in 4% PFA for 15 min, washed 3 times with PBS and then treated with ice-cold MeOH for 5 min on RT. The staining was then performed as described below. Quantification was performed with the Fiji plugin TrackMate [47, 48] to automatically detect puncti with a diameter of 1, 2, and 3 microns at a threshold of 600, 200 and 150, respectively. Nuclei were detected at an average size of 15 microns with a threshold of 10. Unique particles were annotated for 5 different frames for each condition and shown as particles per cell. The experiment was repeated 3 times.

### LD staining and RF microscopy

Samples were plated as for the LC3 staining. For samples treated with oleic acid, 200 µM (Sigma-Aldrich, # O1008-1G) were added at plating, over-night. The next day, samples were incubated in 5 µM BODIPY 493/503 (Invitrogen, #D3922) in PBS and subsequently washed 3 times with PBS and fixed in 4 % PFA for 15 min. Samples were then washed, stained with Dapi for 15 min at RT and mounted with Mowiol. Quantification was performed using the Fiji plugin TrackMate [47, 48] detecting droplets of 1, 2, and 3 microns in diameter at a threshold of 900. Nuclei were detected at 10 micron and a threshold of 1.

For RF microscopy, 250k cells were plated per well of a 6 well glass bottom plate (Cellvis, # P06-1.5H-N) in N2B27 media. After 36 h samples were treated with 200 µM oleic acid o/n and then with BODIPY for 3 h. Samples were washed 3 times with PBS and maintained in N2B27 for RF microscopy. The imaging was performed on Tomocube HT-X1 instrument at 37C and 5 % CO2. At least 5-8 images were acquired for each condition for each well. The z-stacks were acquired with 1.06 µm slices. Max Intensity projections (MIP) are generated on the Tomostudio 3.3.9. The fluorescence max intensity projections were resized to Refractive index tomograms for comparison. The ROIs from each image were isolated for further analysis. Comdet plugin was used for identifying high refractive index regions. The approximate particle size was 2 pixels and Intensity threshold values were chosen individually for each image to avoid any false positives or negatives.

### Immunofluorescence

Samples were washed 3 times with PBS, fixed in 4% PFA for 10 min and again washed 3 times with PBS. Samples stained for LC3 were then treated for 5 min with 100% MeOH at RT and washed again 3 times with PBS. Samples were then blocked and permeabilized with 10% donkey serum (Jackson Immunoresearch, #017-000-121) and 0,1% Triton X (Thermo Fischer, # HFH10) in PBS for 20 min and incubated with primary antibodies at 4 °C over-night. Oct4 (1:100, Satna Cruz, # sc-5279), Nanog (1:100, Abcam, # ab80892), Sox2 (1:50, R&D Systems #AF2018), Lc3 (1:200, Cell Signalling, # 2775S), Pdi (1:200, Cell Signalling, # 3501S), Rcas1 (1:200, Cell Signalling, # 12290S), Rab7 (1:200, Cell Signaling, # 9367S) Sox1 (1:50, R&D Systems, # AF3369), Sox17 (1:200 R&D Systems, # AF1924-SP), Gata4 (1:100 R&D Systems, # AF1700-SP), Gata6 (1:100 R&D Systems, # AF1700-SP), Dab2 (1:100 BD, # 610464). The next day, samples were washed 3 times with 0.1% Triton X in PBS and then incubated with secondary antibodies and Dapi.

### Western Blot Analysis

Cells were lysed in membrane buffer (150 mM NaCl, 50 mM Tris-HCl pH 7,5, 1 mM 2- mercaptoethanol (Sigma, #M3148) and 1% CHAPS (Sigma, #C3023) in water). Lysates were run on SDS-Page and transferred to nitrocellulose membranes. Bands were detected with the Amersham ECL Prime Western Blotting Detection Reagents (#RPN2232) and captured on Fujifilm Super RX-N films (#47410). Antibodies for Vmp1 (Cell Signalling, #12929S), DDIT3 (Cell Signalling, #2895T), β-CATENIN (Santa Cruz, sc-7963), β-ACTIN (BioLegend, W16197A), E-CADHERIN (BioLegend, 67A4), H3 (Abcam, #1791), HA-epitope (Roche, 11867423001), HSP90 (Santa Cruz, #sc- 101494) were used. The Quantification of western blot bands was performed with ImageJ.

### RNA extraction and RT qPCR

RNA was extracted with the RNeasy Kit (Quiagen, #74104) following the manufacturer’s instructions, including the on-column DNA digestion. EB and SCO samples were pre-treated with the QIAshredder (Quiagen, #79656). cDNAs were prepared using the Bio-Rad iScript Reverse Transcription Supermix for RT-qPCR (#1708841). RT qPCR samples were prepared using the KAPPA SYBR R fast (Roche, KK4611). Primer Sequences:

Tmem41b Exon2 FW: TGCAGAAGCTGGATCAGCAA, RV: GAGCCTTGGCGTCATCCATA

Tmem41b Exon7 FW: GCAGGTAGAGCGTCACAGAG, RV: TGCAGAGTTGTTCCTGCCTT

Sdha FW: CGTTATGTGAGGGGTGTGCCTT, RV: CCTTACCAAACCTTGTGTCTGGA

Sox1 FW: ACGTAGCCAACGTAGACAC, RV: ATGAGCGTCGTCCCGTGG

Fgf5 FW: GCCTGTCCTTGCTCTTCCTCAT, RV: GGAGAAGCTGCGACTGGTGA

Actc1 FW: TGTAGGACCGTGTCGAAACC, RV: CGAAACCACACACTGTTACCG

Brachyury FW: TCTCTGGTCTGTGAGCAATGGT, RV: TGCGTCAGTGGTGTGTAATGTG

Gata6 FW: GAAGCGCGTGCCTTCATC, RV: GTAGTGGTTGTGGTGTGACAGTTG

Dab2 FW: TTGATGATGTGCCTGATGCT, RV: TTTGCTTGTGTTGTCCCTGA

Sox17 FW: CCCAACACTCCTCCCAAAGTATC, RV: TTCCCTGTCTTGGTTGATTTCT

Ddit3 FW: GGCTCAAGCAGGAAATCGAG, RV: AAGTGAGAGGCTGTTGACAC

### Quantification and statistical analysis

Statistical analysis was performed using Python and GraphPad Prism. Data are presented as mean-centered and the standard deviation. All experiments were repeated with at least 3 independent biological replicates, unless stated otherwise. The statistical test used to evaluate significance is the Welch’s *t* test. Statistical significance in the figures is denoted as follows: ns: *p*> 0.05, *: *p* < 0.05, **: *p* < 0.01, ***: *p* < 0.001.

### RNA seq

RNA samples were submitted to the Functional Genomics Center in Zurich (FGCZ) for RNA sequencing. The quality of the RNA was determined with a Fragment Analyzer standard sensitivity RNA measurement (SS RNA kit (15 nt), Agilent, Waldbronn, Germany). The measured concentrations (> 50 ng/µl) and RIN (>9.9) values qualified for a Poly-A enrichment strategy in order to generate the sequencing libraries applying the TruSeq mRNA Stranded Library Prep Kit (Illumina, Inc, California, USA). After Poly- A selection using Oligo-dT beads the mRNA was reverse-transcribed into cDNA. The cDNA was fragmented, end-repaired and polyadenylated before ligation of TruSeq UD Indices (IDT, Coralville, Iowa, USA). The quality and quantity of the amplified sequencing libraries were validated using a Fragment Analyzer SS NGS Fragment Kit (1–6000 bp) (Agilent, Waldbronn, Germany). The equimolar pool of samples was spiked into a NovaSeq6000 run targeting 20M reads per sample on a S1 FlowCell (Novaseq S1 Reagent Kit, 100 cycles, Illumina, Inc, California, USA). Reads from RNA-seq were first preprocessed by Trimmomatic (v0.40) [49] to remove low-quality bases and adapters. Then, reads were aligned to mouse genome GRCm38 (release 81 version from Ensembl) using HISAT2 (v2.2.1) [50]. HTSeq Count (v2.0.2) [51] was used to count reads for each gene (Ensembl GRCm38.81), ignoring reads on overlapped regions. Differentially expressed genes were defined by DESeq2 (v1.40.2) [52]. MA plot was generated using custom-made scripts. Genes significantly enriched show a fold change > 1 and FDR (adjusted p-value) < 0.01. The read counts were transformed in RPKM for the heatmap representations of specific genes and pathways. The genes corresponding to germ-layer differentiation markers (including those for embryonic and extra-embryonic endoderm), biological processes, and signaling pathways were obtained through literature mining and the KEGG database (Table S1). The processed data are available as excel files (Table S2 and Table S3) and raw data have been deposited on NCBI SRA database under the accession: PRJNA1031567

### Surface Proteome Profiling

Sample preparation and biotin labelling was performed following the instructions from {Wojdyla, 2020 #140}{Weekes, 2012 #141}. In brief, 10M cells were collected per sample using Accutase (Biolegend, #423201). Cell pellets were washed one with PBS and then incubated, resuspended in 1 mL biotinylation buffer (97 µM aminooxy-biotin, Biotium #90113, 1 mM sodium periodate, Thermo-scientific #20504, 0,093% aniline, Sigma-Aldrich, #242284) for 30 min at 4 °C. Next, cells were pelleted, the reaction quenched with 1 mM glycerol (30min, 4 °C) and washed once with PBS and lysed in 1 mL of lysis buffer (see western blot) by sonication (Dr. Hielscher sonicator, 30 sec at 80% amplitude and 80% cycle time). Lysates were pelleted, the supernatant transferred into a fresh tube and incubated with 30 µL (dry resin) of streptavidin agarose resin (Thermo-Scientific, #20347) for 1.5h at 4 °C. beads collected, transferred onto a column (BioRad, #7326008) and washed with 8 mL of lysis buffer and 4 mL of 0,5% SDS in 50 mM TEAB. Beads were resuspended in 200 µL 50 mM Ambic with 10 mM TCEP (30 min, 37 °C), then 20 mM iodoacetamide was added (30 min. 37 °C). Beads were further washed with 4 mL of SDS buffer (see above), 8 mL Urea buffer (6M Urea in 50 mM TEAB, pH=8,5), and with 2 mL 50 mM TEAB. Beads were taken up in 200 µL TEAB buffer and digested with 3 µL Trypsin (Promega) over night, 37 C. Tryptic digestion was quenched by the addition of acetonitrile (ACN). Tandem mass tag (TMT) isobaric reagents (TMTpro 6plex Thermo Fisher Scientific) were dissolved in anhydrous ACN to a final concentration of 20 mg/ml, of which a unique TMT label was added (100µg). Peptides were incubated at room temperature for one hour, TMT labeling reactions were halted by 0.3% hydroxylamine and dried. Peptides were fractionated by high pH fraction in reverse phase (microspin column, Nest group) following the procedure based on the high pH reversed-phase peptide fractionation kit (Pierce).

LC-MS/MS was performed on an Orbitrap Exploris 480 mass spectrometer (Thermo Fisher) coupled to an Vanquish Neo liquid chromatography system (Thermo Fisher). Peptides were separated using a reverse phase column (75 μm ID x 400 mm New Objective, in-house packed with ReproSil Gold 120 C18, 1.9 μm, Dr. Maisch GmbH) across 180 min linear gradient from 7 to 35% (buffer A: 0.1% [v/v] formic acid; buffer B: 0.1% [v/v] formic acid, 80% [v/v] acetonitrile). Samples were acquired in DDA mode (Data Dependent Acquision) with MS1 scan (scan range = 350-1500, R=120K, max injection time auto and AGC target = 100), followed by dependent MS2 scans (first mass (m/z) 110, R = 45K, max injection time auto and AGC target = 200).Cycle time was set to 2 sec. Peptides with charge between 2-5 were isolated (m/z = 0.7) and fragmented (NCE 36%).

Acquired spectra were searched using the MaxQuant software version 2.1.0.0 against mus musculus proteome reference dataset (http://www.uniprot.org/, downloaded on 03.04.2020) extended with reverse decoy sequences. The search parameters were set to tryptic peptides, maximum two missed cleavage, carbamidomethyl as static peptide modification, oxidation (M) and deamidation (N-terminal) and TMT6 as static peptide modification. The MS and MS/MS mass tolerance was set to 10 ppm. False discovery rate of < 1% was used at PSM and protein level. Abundance of 1417 proteins was determined from the intensity of top two peptides. Intensities values were median normalized and missing values were imputed using random sampling from a normal distribution generated from 1% less intense values. Statistical analysis was performed using unpaired two sided t-test and pvalues were corrected using Benjamin-Hochberg correction. The entire dataset, including raw data, generated tables and scripts used for the data analysis are available in the PRIDE repository: PXD053039. Matrices with protein intensities are reported in Table S4.

## Acknowledgments

We acknowledge the Functional Genomics Center Zurich (FGCZ) for technical support and Sungsik Lee from ScopeM facility for help on the RF microscopy. We thank G.J. Pereira, C. Fimiani, and U. Sutter for their help with reagents. We thank C. Ciaudo, A. Grison, J. Corn, and K. Basler for their helpful discussions. GDM was supported by the ETH Zurich Postdoctoral Fellowship Program as well as the Marie Curie Actions for People COFUND Program. This work was supported by grants from the Swiss National Science Foundation (SNF grants 31003A_152814/1 and 31003A_175643/1).

## Author Contributions

Conceptualization: MH, AW and GDM; Data curation: MH, FU, VG and GDM; Formal analysis: MH, TS, HH, FU, VG, and GDM; Funding acquisition: KW, AW and GDM; Investigation: MH, AW and GDM; Methodology: MH, TS, HH, FU, VG, WJ, and GDM; Project administration: AW, GDM; Resources: MH, KW, AW and GDM; Supervision: KW, AW and GDM; Validation: MH, GDM; Visualization: MH, FU, VG and GDM; Writing – original draft: MH, AW and GDM; Writing – review & editing: MH, TS, HH, FU, VG, WJ, KW, AW and GDM.

