## Supplementary Figures for "The scramblases VMP1 and TMEM41b are required for primitive endoderm specification by targeting WNT signaling"

### SUPLEMENTAL FIGURE LEGENDS

#### S1 Characterization of single and double mutant ESCs

**(A)** Efficient depletion of VMP1 protein in Vmp1 and Double KO ESCs. Western Blot analysis shows VMP1 expression and HSP90 as housekeeping. **(B)** PCR analysis of the deletion induced by CRISPR-Cas9 at the Tmem41b genomic locus in WT and mutant cells. WT cells show amplification of a product of 3000 bps; in Tmem41b and Double KO ESCs, a fragment of 500 bps is detected. The PCR negative control (C-) is indicated. For each mutation, two independent clones were characterized. **(C-D)** RT-qPCR analysis showing decreased Tmem41b expression in Tmem41b and Double KO ESC using exon-spanning primers specific for exon 2 **(C)** and exon 7 **(D)**. mRNA levels are relative to WT ESC sample and normalized to Sdha mRNA. **(E)** Single and Double KO ESCs show similar proliferation rate. Plotted is proliferation factor after 48h. Each dot represents independent experiments. For each KO condition, two independent clones were tested. **(F-G)** Single and double KO ESCS show similar expression of pluripotency markers. **(F)** Representative IF images showing WT and mutant ESCs stained for the pluripotency markers NANOG, OCT4, and SOX2. Nuclei are stained with Dapi. Scale bar: 20 µm. **(G)** Quantification of pluripotency markers from **F** relative to the total cell number. Each dot represents an independent frame analyzed. **(H)** Morphology of single and double KO ESCs is like WT ESCs. Brightfield images showing ESCs morphology. Scale bar: 100 µm. **(I-J)** Transcriptomic analysis of VMP1 **(I)** and TMEM41b **(J)** KO ESCs showed only limited transcriptional differences, plotted here as MA plot. Genes differentially expressed are labeled in blue. Genes highlighted in red are pluripotency markers.

#### S2 Characterization of Autophagy and Lipid Droplet phenotypes

**(A)** Single and Double KO ESCs show accumulation of LC3+ autophagosomes. Representative IF images show ESCs stained for LC3; nuclei are stained with Dapi. Scale bar: 20 µm. **(B)** Single and Double KO ESCs show increased number of autophagosomes per cell. Graph shows quantification of LC3 puncta per cell of (A) in WT, Vmp1, Tmem41b, and double KO ESCs. Each dot represents an analyzed frame. Two independent mutants were analyzed per KO condition. **(C)** Autophagosomes in double KO ESCs are bigger than in single KO conditions. Pie charts show the size distribution of LC3 puncta of 1, 2, and 3 µm diameter in each mutant condition. Two independent mutants were analyzed per mutant condition. **(D)** Double KO ESCs accumulate LD. Live cell staining of WT, Vmp1, Tmem41b, and double KO ESCs with BODIPY. Cells were pre-treated with or without oleic acid (OA). **(E)** Quantification of LD frequency per cell in the absence or in the presence of OA treatment. **(F)** LDs are visible by Refractive index microscopy. Representative image of Double KO ESCs treated with OA highlighting a calibration bar for refractive index. White spots correspond to a refractive index of 1.4, which is specific for LDs. Scale bar: 20 µm. **(G)** LDs in refractive index microscopy overlay with BODIPY staining. Representative images show RF images co-stained with BODIPY (bottom panel). Scale bar: 10 µm.

**S3 Vmp1, Tmem41b, and Double KO ESCs show similar morphology and transcriptional profile**

**(A)** Single and double KO ESCs do not show changed morphology of intracellular organelles. Representative confocal IF shows ESCs stained for RCAS1 (Golgi, left), RAB7 (late endosomes, center), and PDI (ER, right) in green; nuclei were stained with Dapi. Scale bar: 20 µm. **(B-E)** Transcriptional analyses of WT and Vmp1, Tmem41b, and Double KO ESC show no significantly deregulated genes. **(B)** PCA analysis of the transcriptional profile of WT, single, and Double KO ESCs. **(C)** Diagram showing uniquely and collectively differentially expressed genes in Vmp1, Tmem41b, and Double KO ESCs. **(D)** Differentially expressed genes in single and Double KO ESCs are not enriched for specific developmental processes. The plot illustrates the differentially expressed genes based on germ layers, with the size of dots indicating the amount of differentially expressed genes. **(E)** Differentially expressed genes in single and double KO ESCs are not enriched for specific signaling pathways. The plot illustrates the differentially expressed genes based on molecular pathways, with the size of dots indicating the amount of differentially expressed genes.

**S4 Vmp1, Tmem41b and Double KO ESCs differentiate into the three germ layers, but not into the Dab2 positive primitive endoderm**

**(A)** ESCs differentiated to EBs express lineage markers for ectoderm (Fgf5), mesoderm (Brachyury). They show decreased expression of the pluripotent marker Pouf51. Graphs show mRNA levels of WT ESCs and EBs normalized to the Sdha gene. Each dot represents an independent experiment. **(B)** RT-qPCR analysis for Fgf5 and Brachyury mRNA expression in mutant EBs relative to WT EBs. Two independent clones were analyzed per cell line. Each dot represents an independent experiment. The line at 0,7 indicates an arbitrary threshold after which successful expression of the marker is considered. **(C)** The heatmap summarizes the expression of a given marker across all experiments and compares it to the WT EBs expression profile by calculating p values. (**D)** Double KO ESCs differentiate into SOX1 expressing Spinal Cord organoids (SCO). Representative IF images of SCOs stained for the neuronal stemness marker SOX1; nuclei are stained with Dapi. Scale bar: 50 µm.

**S5 Double KO ESCs show a delay in XEN cell specification**

**(A-B)** Double KO ESCs differentiate at low efficiency into XEN cells. **(A)** Graphs show mRNA levels of WT and Double KO XEN at day 6. Data were normalized to the Sdha gene and expressed relative to mRNA level detected in WT ESCs. Each dot represents an independent experiment. **(B)** Representative IF images of day 6 XEN cells stained for the XEN markers DAB2 and GATA6 in the left panel and GATA4 and SOX17 in the right panel; nuclei are stained with Dapi. Scale bar: 150 µm. **(C)** XEN cells derived from Double KO ESCs can be maintained in culture. Brightfield images show WT and Double KO XEN cells at passage 3. Scale bar: 50 µm **(D-F)** Transcriptional analysis of XEN cells. **(D)** Differentially expressed genes in Double KO vs. WT conditions do not cluster by biological processes. Plot shows differentially expressed genes over biological processes. The size of dots corresponds to the number of differentially expressed genes. **(E)** Double KO XEN cells show decreased expression of XEN markers. Heatmap comparing min to max expression of XEN markers in WT and double KO XEN cells.

**S6 Analysis of protein abundance at the plasma membrane in WT and VMP1$^{KO}$ cells.**

**(A)** Heatmap depicting sample correlation in the MS analysis. **(B)** GO term analysis comparing protein enrichment between plasma membrane-purified samples and whole-cell lysates. **(C)** Volcano plot illustrating differentially expressed proteins at the plasma membrane in WT and VMP1KO ESCs. Proteins with the most significant differences are highlighted in red. Gray lines indicate the selection criteria: a log2 fold-change > 1 and a false discovery rate < 0.01. The positions of Frizzled proteins are annotated. **(D)** mRNA expression levels of Frizzled genes in WT and VMP1KO ESCs, as shown in the transcriptomic profile described in Fig. 1.

**S7 WNT signaling upregulation rescues the defect in XEN specification of Double KO cells.**

**(A-B)** Chiron-dependent WNT activation rescues the delay of Double KO XEN differentiation. **(A)** Brightfield images show XEN clusters of Double KO XEN cells (highlighted by white lines) with and w/o Chiron treatment. Scale bar: 100 µm. IF staining for DAB2 and GATA6 **(B)**; nuclei are stained with Dapi. Scale Bar: 150 µm. **(C)** Derivation of ESC clones with HA-FZD2 over-expression. Western blot showing HA-FZD2 expression in WT and Double KO cells. ACTIN is used as a loading control. **(D-F)** FZD2 overexpression rescues the differentiation delay in Double KO XEN cells. Immunofluorescence staining for SOX17 and GATA4 **(D)** or DAB2 and GATA6 **(E)** in WT and Double KO cells. Nuclei are counterstained with DAPI. Scale bar: 100 µm. (D) Quantification of GATA6 and DAB2 positive cells during XEN differentiation, expressed as a percentage of total cells (DAPI-stained).
